# Cold and distant: structural features of the nucleoprotein complex of a cold-adapted influenza A virus strain

**DOI:** 10.1101/847608

**Authors:** A.V. Shvetsov, D.V. Lebedev, Y.A. Zabrodskaya, A.A. Shaldzhyan, M.A. Egorova, D.S. Vinogradova, A.L. Konevega, A.N. Gorshkov, E.S. Ramsay, A. Radulescu, M.V. Sergeeva, M.A. Plotnikova, A.B. Komissarov, A.S. Taraskin, K.I. Lebedev, Yu.P. Garmay, V.V. Kuznetsov, V.V. Isaev-Ivanov, A.V. Vasin, L.M. Tsybalova, V.V. Egorov

## Abstract

Two influenza A nucleoprotein variants (wt: G102R; and mutant: G102R and E292G) were studied with regard to macro-molecular interactions in oligomeric form (24-mers). The E292G mutation has been previously shown to provide cold adaptation. Molecular dynamics simulations of these complexes and trajectory analysis showed that the most significant difference between the obtained models was distance differences between nucleoprotein complex strands. Influenza virus nucleoprotein complexes were isolated from strains bearing the corresponding NP amino acid substitutions. The isolated complexes were characterized by transmission electron microscopy and differential scanning fluorimetry (DSF). Presence of the E292G substitution was shown by DSF to affect nucleoprotein complex melting temperature. In the filament interface peptide model, it was shown that the peptide corresponding in primary structure to the wild-type NP (SGYDF**<u>E</u>**REGYS, wild type peptide) is prone to temperature-dependent self-association, unlike the peptide carrying the substitution corresponding to E292G (SGYDF**<u>G</u>**REGYS, mutant peptide). It was also shown that the SGYDF**<u>E</u>**REGYS peptide (wt) is capable of interacting with a recombinant full-size monomeric nucleoprotein (with primary structure corresponding to wild type); this interaction’s equilibrium dissociation constant is five orders of magnitude lower than for the SGYDF**<u>G</u>**REGYS peptide. Using small-angle neutron scattering (SANS), the supramolecular structures of isolated complexes of these proteins was studied at temperatures of 15, 32, and 37°C. SANS data show that the structures of the studied complexes (mutant or normal proteins with RNA) at elevated temperature differ from the rod-like particle model and react differently to temperature changes. The data suggest that the mechanism behind cold adaptation with E292G is associated with a weakening of the interaction between strands of the ribonucleoprotein complex and, as a result, the appearance of inter-chain interface flexibility necessary for complex function at low temperature.

## Introduction

The incidence of influenza is detrimental to human health and is a serious economic concern. Vaccination is one of the most effective ways to prevent the occurrence of this illness [1]. It is advisable to vaccinate using live attenuated vaccines [2], including vaccines created using cold-adapted, temperature-sensitive strains as donors of attenuation [3]. The NP protein is one of eight major structural proteins of the influenza virus [4]. As a ribonucleoprotein particle component, it takes part in both the packaging of viral genetic material and in replication [5]. In the vaccine strain obtained during cold adaptation of the influenza A/HK/68 virus, a mutation leading to the amino acid substitution E292G in the NP protein was detected [3]. This substitution is sufficient for the virus to acquire a cold adaptation phenotype. The presence of another substitution (G102R), however, is not related to the acquisition of this property directly, but is necessary for the virus to multiply when the E292G substitution is present [6]. The molecular mechanisms behind the acquisition of cold adaptation, through these substitutions, remain unclear. Elucidation of such mechanisms is necessary for rational design of new strains, including donors of attenuation necessary for the creation of new live attenuated vaccines. The presence of certain substitutions in NP (G102R, E292G) leads to cold adaptation, but not temperature sensitivity. E292G was shown [6] *in vitro* to be insufficient to confer temperature sensitivity. Moreover, E292G is also seen *in vivo.* It has been detected in viruses infecting wild birds (whose body temperatures are commonly 39°C and above).

In this research, we study the influence of the substitutions G102R and E292G on ribonucleoprotein complex structure using molecular dynamics, differential scanning fluorimetry, and small-angle neutron scattering.

## Materials and methods

### Molecular Modeling

To build NP filaments models, we used existing cryoelectron microscopy and initial model data available from pdbid:4bbl [5]. Full monomer models were built using the Modeler software package and pdbid:2iqh [7] as template. Initial structure was modified as follows: wild-type NP contains the substitution G102R; and mutant (cold-adapted) NP contains two substitutions simultaneously (G102R and E292G). Next, each monomer was fitted to the 4bbl structure. Then, RNA was added using P atoms trace in 4bbl. RNA was built using the Ambertools package. The resulting structure was then used for iterative optimization using Pymol [8] and GROMACS [9] to resolve all clashes.

### Molecular dynamics

All MD simulations were performed in the GROMACS [9], [10] software package using an amber99sb-ildn-bsc0 [11], [12] force field for protein and RNA, and a tip3p [13] model for explicit water. The simulated systems were around 2.6M atoms in size, including an explicit solvent shell and 50 mM NaCl to neutralize charges. Solvent shell thickness was at least 2.0 nm. Box size was around 50×22×24 nm. Systems were simulated at two temperatures, 299K and 312K. For each temperature, systems were equilibrated using a two-step protocol. During the first step, the system was equilibrated for 5 ns with all heavy, non-solvent atoms restrained to their initial positions using NPT ensemble. During the second step, systems were equilibrated without restraints for 10 ns (starting from the last frame of the previous step). After two equilibration stages, a 250 ns trajectory was simulated with a time step of 2 fs. A Neighbor search was performed every 50 steps. The PME algorithm was used for modeling electrostatic and Van der Waals interactions with a cutoff of 1.2 nm. Temperature coupling was done with the Nose-Hoover algorithm. Pressure coupling was done with the Parrinello-Rahman algorithm for 1 bar. All further analyses were performed on the final 100 ns time interval.

### RNPs

Isolation of influenza A H3N2 virus (A/Hong Kong/68) ribonucleoprotein complexes was carried out using a viral suspension kindly provided by the Smorodintsev Research Institute of Influenza’s Influenza Vaccine Laboratory, following published methods [14]. Identification of the NP-containing fractions was carried out using described Western blotting methods [15]. For SANS, the protein was transferred to a heavy water buffer. Two hundred microliters of mix (combined, NP-containing pooled glycerol fraction mix, purity above 95%) was applied to a micro-centrifuge filter (with a pore size corresponding to a molecular weight of 100 kDa) and centrifuged until the volume was reduced to 50 μl. Next, 150 μl of PBS (in D20) was added and the centrifugation wash procedure repeated 3 times.

### TEM

For transmission electron microscopy, a suspension of the isolated nucleoprotein complexes at a concentration of 1 mg/ml was deposited on copper electron microscopy grids coated with a carbon substrate. After adsorption of RNP complexes to the substrate for 1 min, the grids were washed twice with distilled water. Further, negative contrasting of the samples on grids with a 2% solution of sodium salt of phosphoric tungsten acid was performed for 1 min. After contrasting, grids were dried and examined using a JEOL JEM 1011 transmission electron microscope at an accelerating voltage of 80 kV. Electron micrographs were taken using a Morada digital camera (Olympus, Inc). The procedure was repeated 5 times.

### DSF

Thermal protein denaturation stability analysis was performed via differential scanning fluorimetry (using intrinsic fluorescence of tryptophan residues) by slowly heating the sample in a Prometheus NT.48 scanning fluorimeter (NanoTemper Technologies, GmbH). Nucleoprotein samples in PBS buffer solution (at concentrations of 3 mg/ml) were loaded into quartz capillaries (Standard Capillaries Prometheus NT.48, NanoTemper); capillaries were placed on the platform and measurements were taken in the temperature range from 15°C to 95°C in increments of 1°C /min at a laser intensity of 50%. Fluorescence intensities were recorded at 330 nm and 350 nm; and the ratio of the fluorescence intensities and the first derivative were analyzed.

### SPR

Measurements were taken using a GE Healthcare Biacore X100 instrument in accordance with the manufacturer’s instructions. Commercial NP protein (H3N2) was covalently immobilized on CM5 chips (GE Healthcare, USA) and used as a ligand. As a result of optimization of conditions, NP immobilization was carried out in a 10 mM MES buffer, pH 6.0. Solutions of the SGYDF**<u>E</u>**REGYS and SGYDF**<u>G</u>**REGYS peptides in PBST buffer (GE Healthcare, USA) at the indicated concentrations were used as analyte. Data were processed in the Biacore X100 software and visualized in Origin2015.

### SANS

Small-angle neutron scattering spectra of the purified normal and mutant NP particles were measured at 15, 32 and 37°C. Measurements were performed on a KWS-1 spectrometer (JCNS at MLZ, FRM-2 reactor, Garching, Germany) at three separate detector positions covering the *q* range from 0.003 to 0.25 Å^−1^. Gyration radius *Rg* and *P(R)* function were calculated using ATSAS software (PRIMUSQT [16] and GNOM [17], respectively). Origin2015 software was used for visualization.

## Results and discussion

In order to examine the role of the protein region containing the mutated residue in cold adaptation, we performed molecular dynamics simulations based on structures obtained by modification of a published structure (4BBL) [5]. In that published structure, it is clear that the amino acid residue that underwent substitution is located at the interface between double protein helix strands. We performed molecular dynamics of the mutant and normal proteins at two temperatures: 26°C, the known lowest tolerable replication temperature for the mutant; and 39°C, the normal temperature for virus propagation. Figure 1 shows several aspects of NP structure: the left panel shows the representative structure obtained from the published RNP structural data [5]; the center panel details structures obtained as a result of molecular modeling, built into the same electron density distribution from the published RNP structure; and the right panel shows the position of amino acid residue 292E, which is substituted by G in NP from the cold-adapted strain.

**Figure 1.**
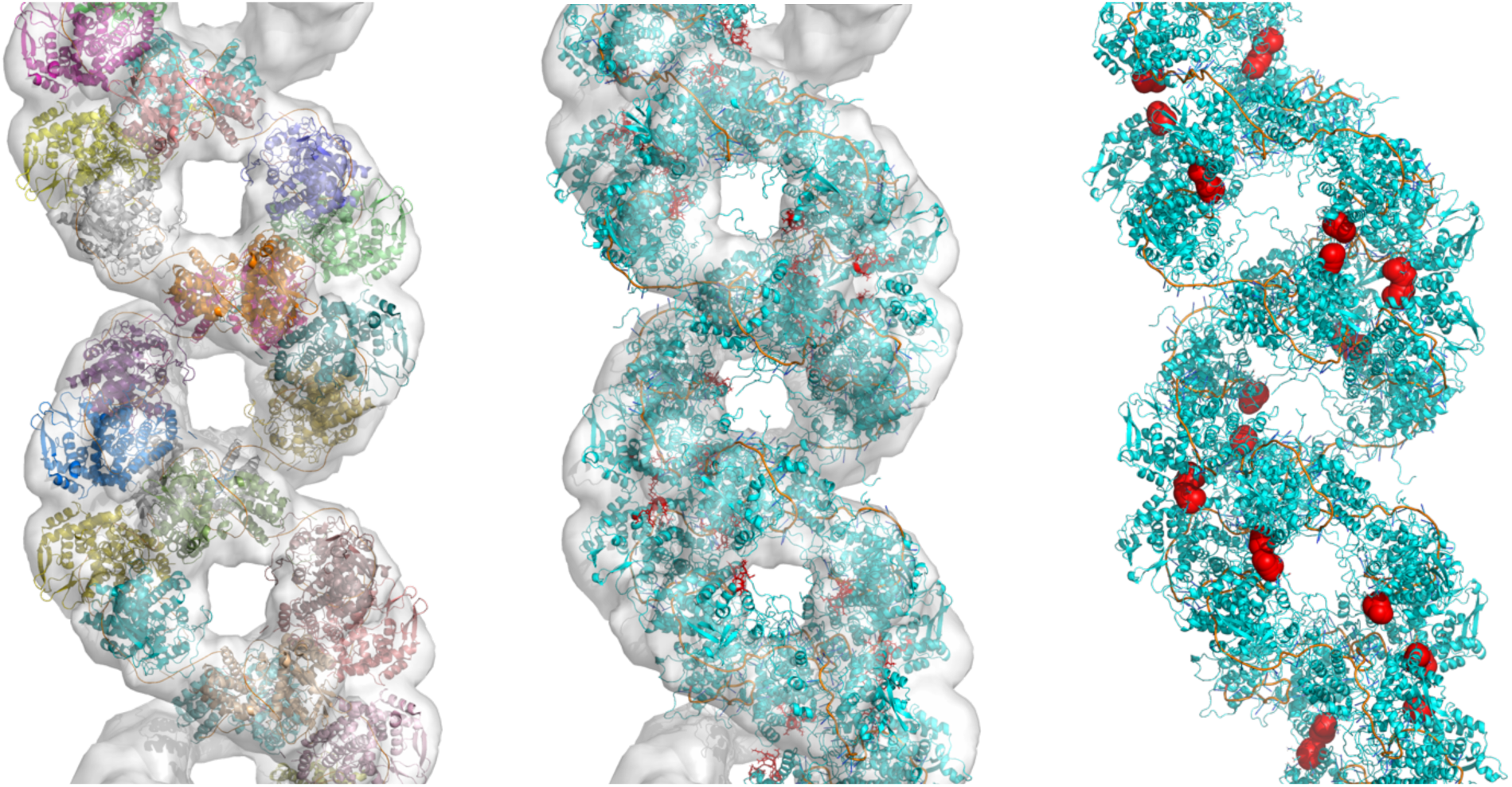
Several details of published and modeled NP structure (discussed in text).

According to the published RNP structure (the starting point for modeling dynamics), identical protein strand regions near position 292 are located opposite each other (Figure 1, right panel). Figure 2 shows an enlarged section of the ^287^SGYDF**<u>E</u>**REGYS^297^ protein fragment located near residue 292.

**Figure 2.**
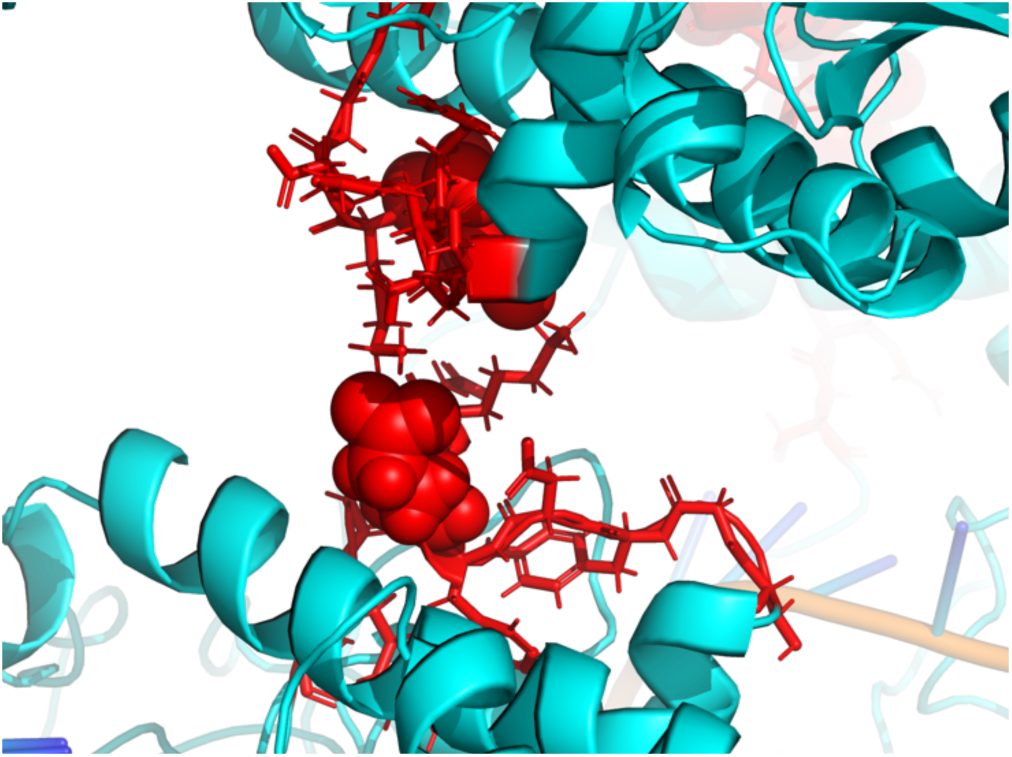
Enlarged rendering of the interface between ribonucleoprotein chains. The ^287^SGYDF**<u>E</u>**REGYS^297^ fragment is shown in red; the amino acid residue 292 (E) is shown in ball representation.

The distances between 292 residues of the adjacent monomers within the same chain, obtained by an analysis of four replicate MD trajectories, exhibited a large degree of scatter due to the filament structure’s flexibility (Figure 3). While, on average, these distances were slightly shorter than those observed in the published RNP structure (63 Å, denoted by horizontal lines in plots), these results show no evidence of steric interactions or any strong allosteric coupling in this set of monomer pairs. The distances do not show temperature dependency, nor do they undergo any significant changes in the mutant protein, as compared to wt (Figure 3).

**Figure 3.**
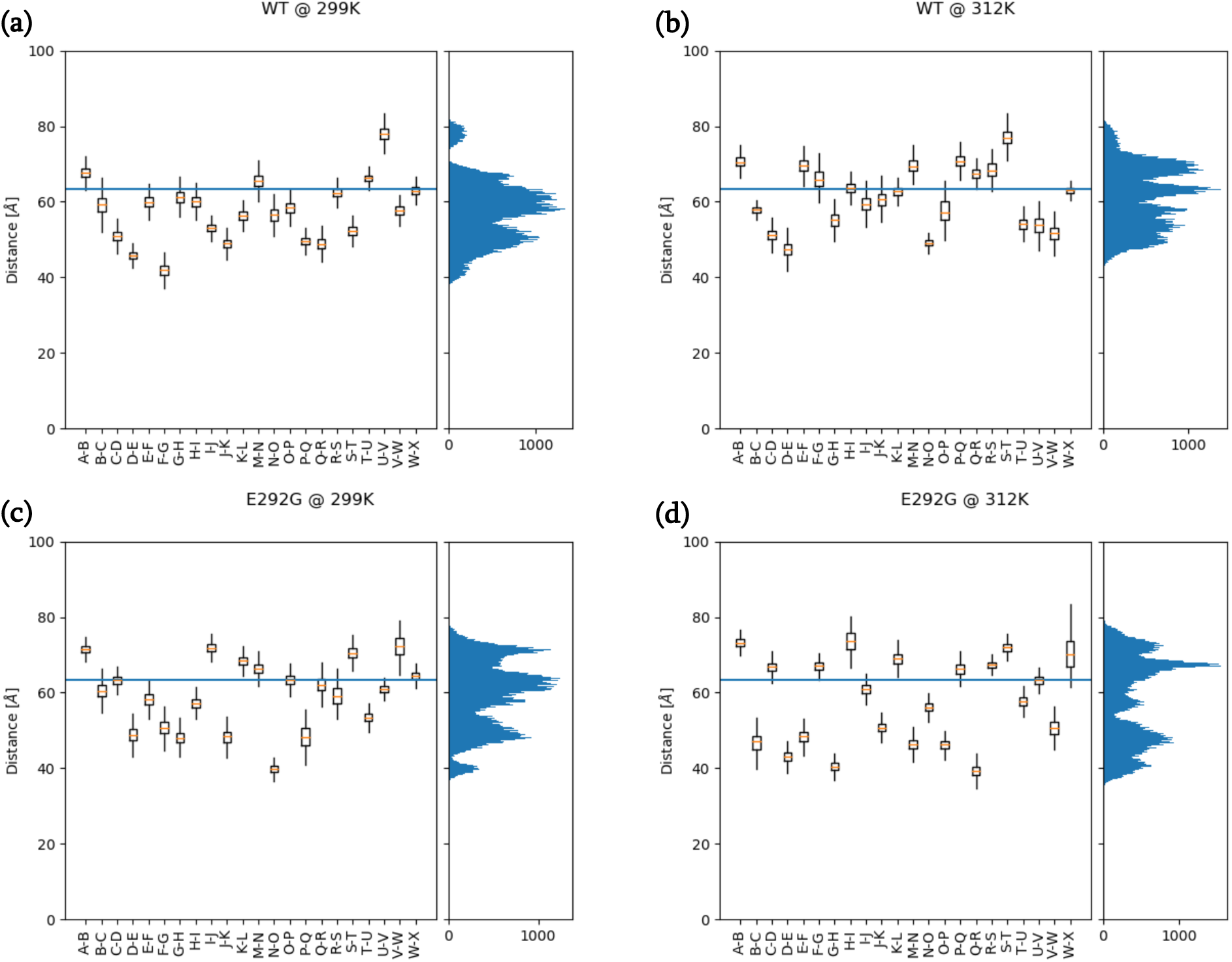
Distances between 292 residues within the same chain of wild-type (wt) and mutant (E292G) RNP at two temperatures: 26°C (299K) and 39°C (312K). The histograms on the right show the detailed distribution of distances (Counts vs Distance).

Analysis of MD data on distances between the 292 residues of monomers located in opposing chains indicates a strong interaction between these regions in both wt and mutant protein. However, while the published structure implies an ordered filament with the distances in question of approximately 10 Å, only about half of these pairs maintained contact throughout our MD simulations (Figure 4). Raising the temperature from 26 to 39°C appears to stabilize some of the contacts, seen as decreased mean distance between the interacting residues. This may be mediated by alleviation of geometric constraints via increased large-scale filament flexibility at the higher temperature and emergence of pairs with very large distances (> 40 Å). The mutant protein filament was characterized by mean distances between 292 residues (on opposing chains) that were larger than in the wild type, as well as by the presence of pairs with distances over 40 Å at both 26°C and 39°C.

**Figure 4.**
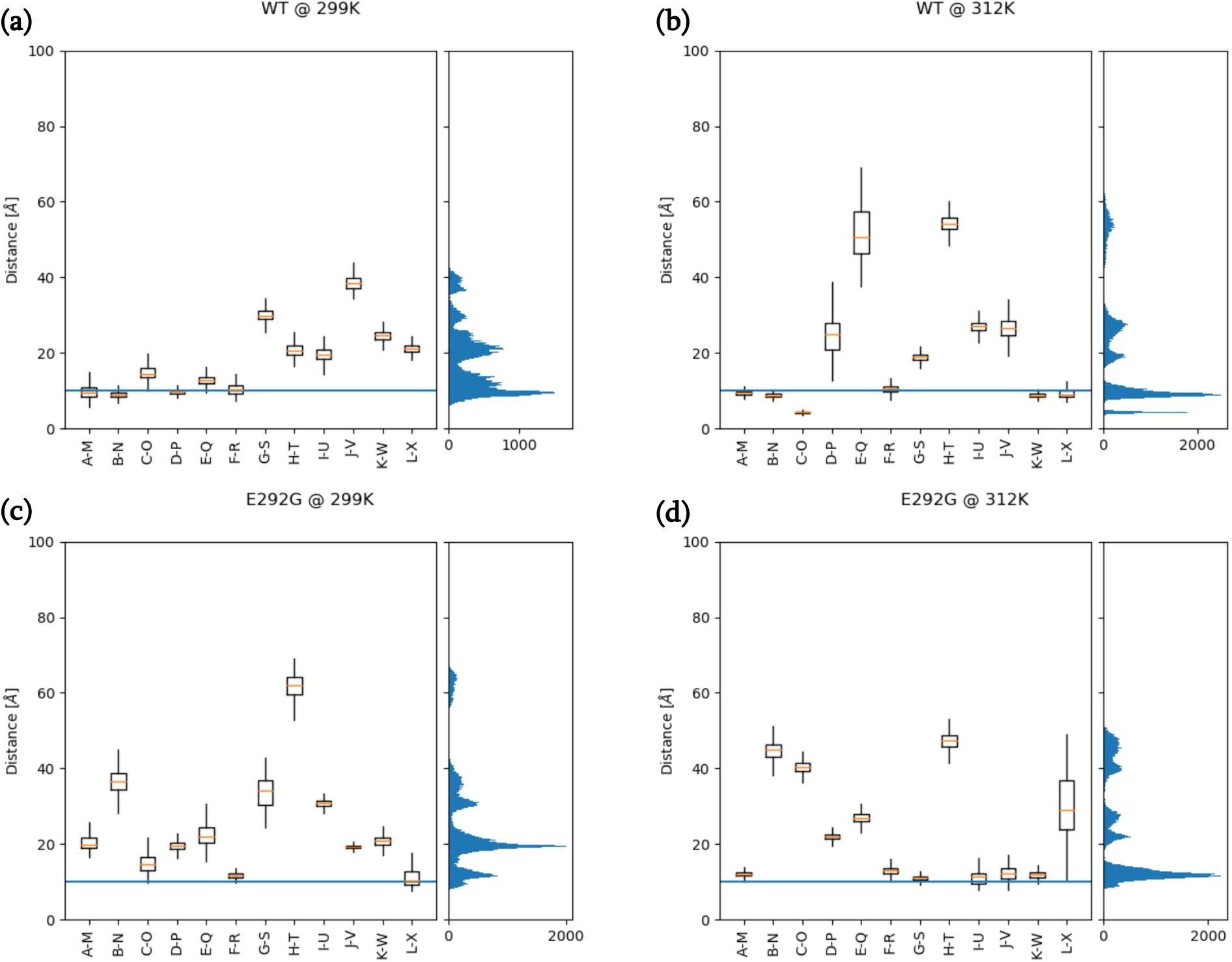
Distances between 292 residues in opposing chains of wild-type (wt) and mutant (E292G) RNP at two temperatures: 26°C (299K) and 39°C (312K). The histograms on the right show the detailed distribution of distances (Counts vs Distance).

The detected increase in the distance variability between the double-stranded protein helix chains (at a lower temperature in the protein carrying the E292G substitution) allowed us to hypothesize that the cold adaptation phenotype depends on this structural feature of NP filaments. Functionally, this can be explained by known influenza A RNA-dependent RNA polymerase mechanisms. Specifically, regulation of interaction between protein strands in RNPs is critical for the operation of the enzyme [5]. In addition, the ability of the double protein helix to unwind and other significant changes in RNP helical structure have been shown [18]. We have previously published differences in RNP structure in solution at room temperature (22°C) and operating temperature (37°C), according to small-angle X-ray scattering data [15]. It is important to note that in different strains featuring different reproduction temperature characteristics, various molecular mechanisms may be behind the cold-adapted phenotype. A detailed review of the currently known cold-adapted or temperature-sensitive strains, including information on amino acid substitutions that affect the ribonucleoprotein complex or temperature features, has been published [19]. In order to verify properties predicted by computer modeling, RNPs from mutant and the wild-type influenza strains were isolated, purified, and characterized. Isolations were carried out according to described methods [14] using centrifugation in glycerol density gradients. Electron micrographs of the obtained RNPs, formed by normal and mutant proteins, are shown in Figure 5.

**Figure 5.**
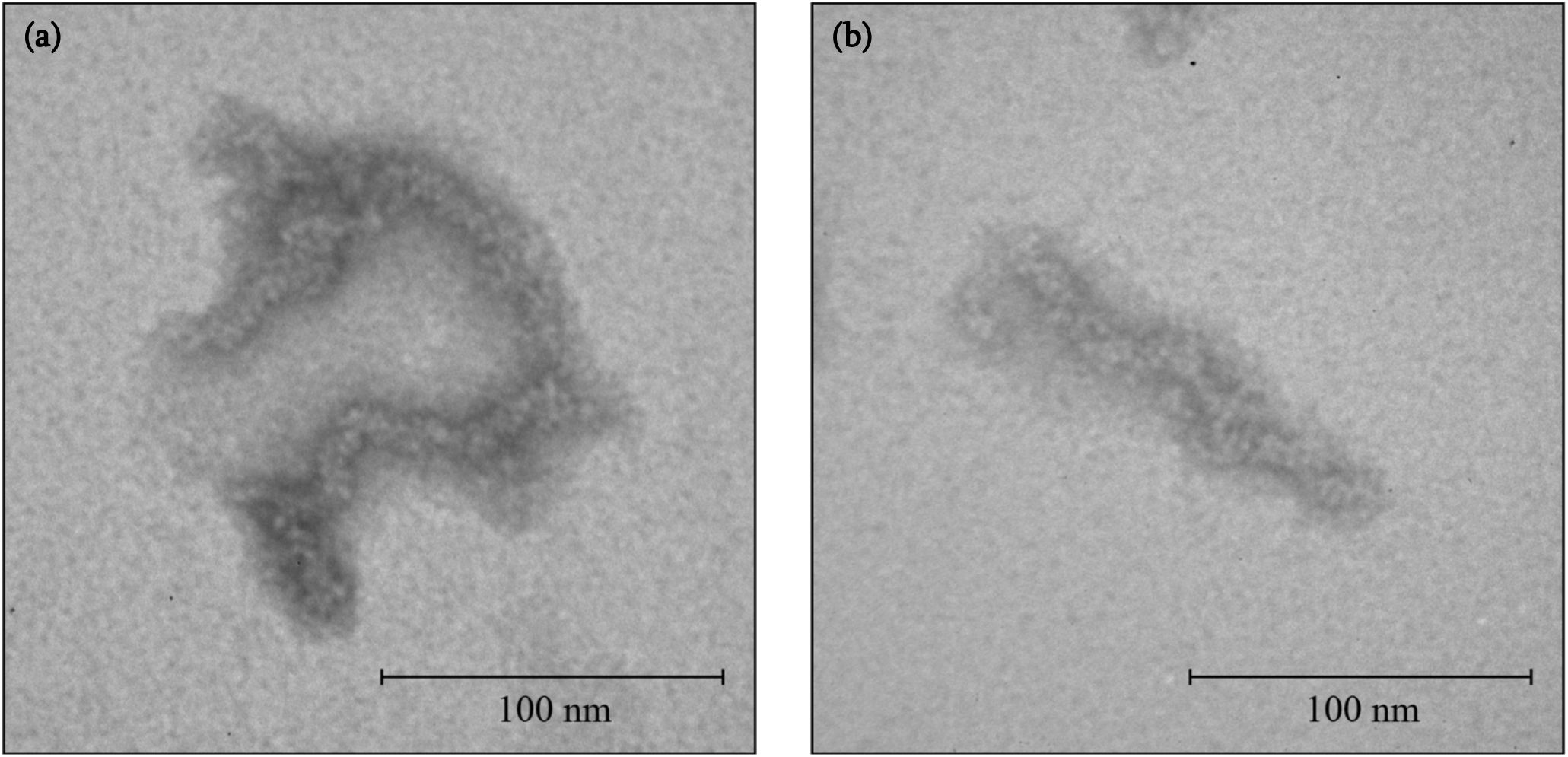
Electron microscopy of ribonucleoproteins isolated from normal (left) and mutant (right) protein-carrying strains.

Differential scanning fluorimetry can be used to support findings collected using molecular dynamics (temperature-dependent changes in RNP structure). Briefly, a change in the 350/330 nm tryptophan self-fluorescence intensity ratio is observed over a temperature gradient from 15 to 95°C. It signifies a change in the tryptophan micro-environment, as dependent on temperature [20]. The experimental results are shown in Figure 6.

**Figure 6.**
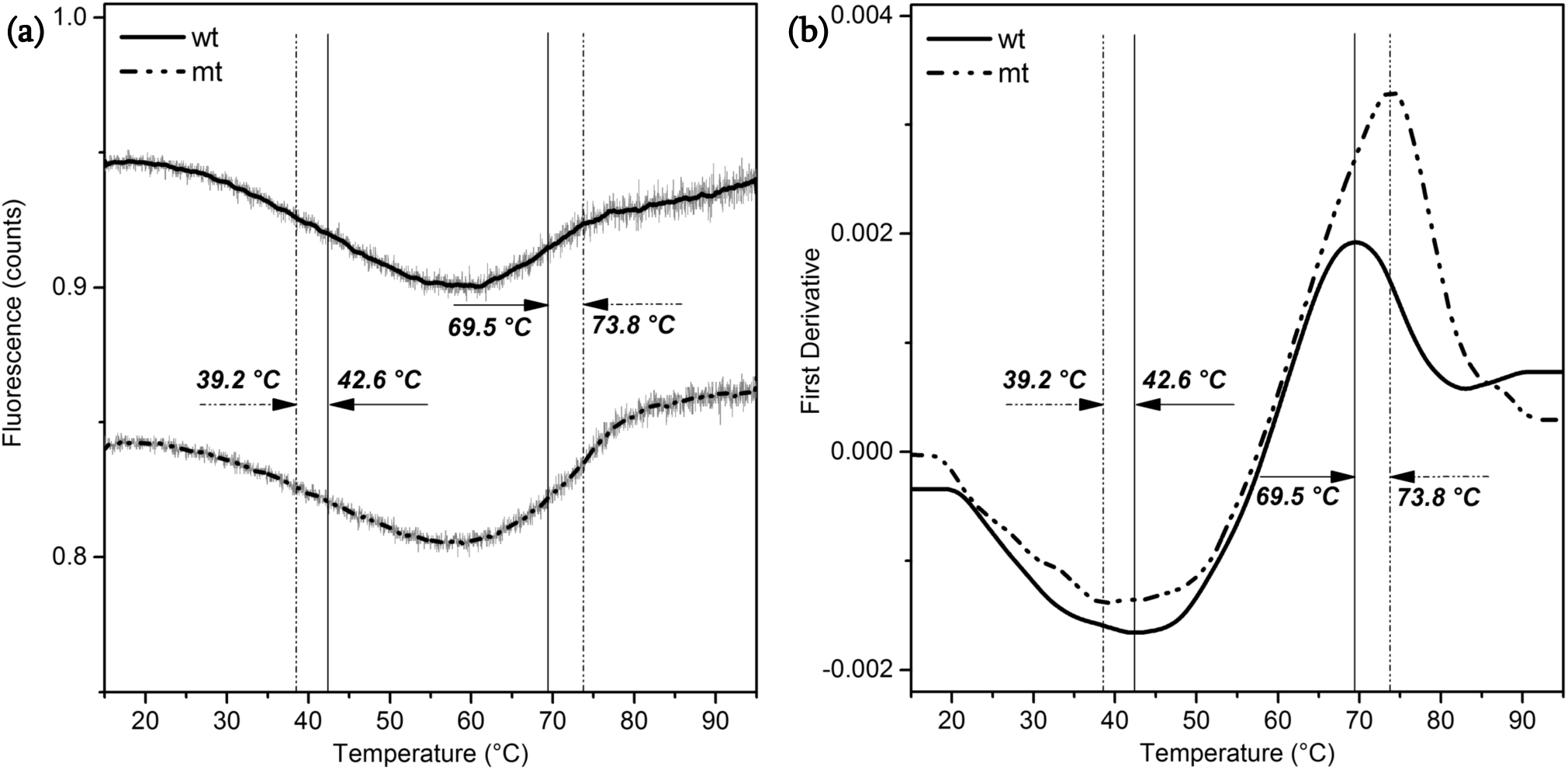
Differential scanning fluorimetry (DSF): (a) intrinsic fluorescence (350/330 nm ratio) of tryptophan as a function of temperature in samples containing wild-type (wt) or E292G mutant (mt) RNP; (b) first derivative of (a).

Analysis of the first derivative of the 350/330 fluorescence intensity ratio shows the presence of a peak characteristic of NP (about 70°C) [21]. Additional peaks, differing from each other, and corresponding to the local melting points of wild type and mutant protein-containing RNPs (42.6°C and 39.2°C, respectively) were seen that can be attributed to partial double protein helix unwinding. The obtained results indicate an influence of the studied mutation on the temperature dependence of the RNP structure. However, the data do not permit pinpointing of the exact RNP structural element responsible for the changes observed. We assert that the changes in the distance between position 292 residues (between nucleoproteins of opposing RNP chains), observed during simulation, are related to the temperature dependence of the interaction at the corresponding interfaces (shown in Figure 2). To test the hypothesis that interactions at the interface play a role, not allosteric effects acting on the chains from elsewhere, we applied an interface peptide model. A commercial NP protein from influenza H3N2, which does not carry the E292G substitution, was immobilized onto a chip. The interaction between chip and either of two peptides (SGYDF**<u>E</u>**REGYS or SGYDF**<u>G</u>**REGYS), which are peptide analogues of the RNP protein strand interaction interface, was studied by surface plasmon resonance (Figure 7).

**Figure 7.**
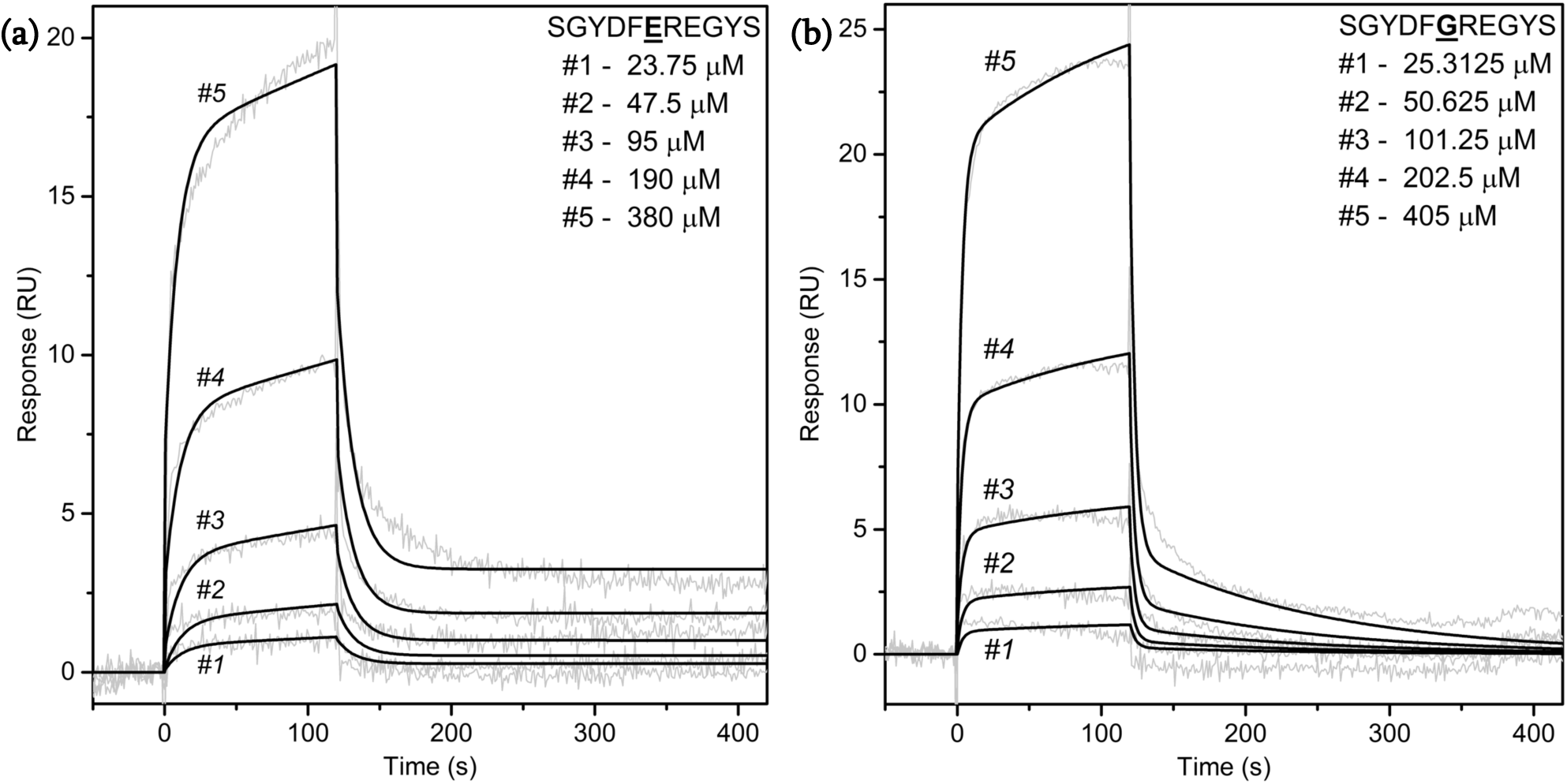
Sensograms of the interaction of peptides, whose primary structures correspond to wt and mutant (E292G) NP protein interfaces, with monomeric NP immobilized on chip; surface plasmon resonance data.

The macromolecular interaction data are best modeled mathematically (Table 1) using a two-step reaction model [22]:

**Table 1.** Constants determined by the two-step reaction model.

| | $k_{a1}$ (1/Ms) | $k_{d1}$ (1/s) | $k_{a2}$ (1/s) | $k_{d2}$ (1/s) | $K_D$ (M) |
| --- | --- | --- | --- | --- | --- |
| SGYDFEREGYS | 71.48 | 0.0799 | 0.0027 | **1.54E-06** | 6.32E-07 |
| SGYDFGREGYS | 17.09 | 0.2899 | 0.0032 | **0.0076** | 0.0115 |

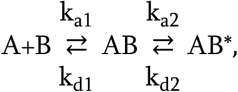

where k_a_ – association rate constant, k_d_ – dissociation rate constant, K_D_ – equilibrium dissociation constant.

The only significant difference between the study peptides, in terms of this model, was the reverse reaction constant k_d2_ (AB* to AB). Specifically, in the case of the SGYDF**<u>E</u>**REGYS (wild-type) peptide, this reverse transformation is slow. However, with the SGYDF**<u>G</u>**REGYS (E292G mutant) peptide, the forward and backward reactions rates are comparable. Thus, it turns out that the unsubstituted peptide is more prone to interaction with the full-sized immobilized protein. At a fixed temperature (25°C), the interactions between the immobilized protein and peptides from solution were different, which may be due to the role of E at position 292 in interface interactions. We also performed an analysis of study peptides in terms of temperature-dependent transition from monomeric to oligomeric (aggregated) form. Figure 8 shows the temperature dependence of 540 nm absorption for the SGYDF**<u>E</u>**REGYS and SGYDF**<u>G</u>**REGYS peptides.

**Figure 8.**
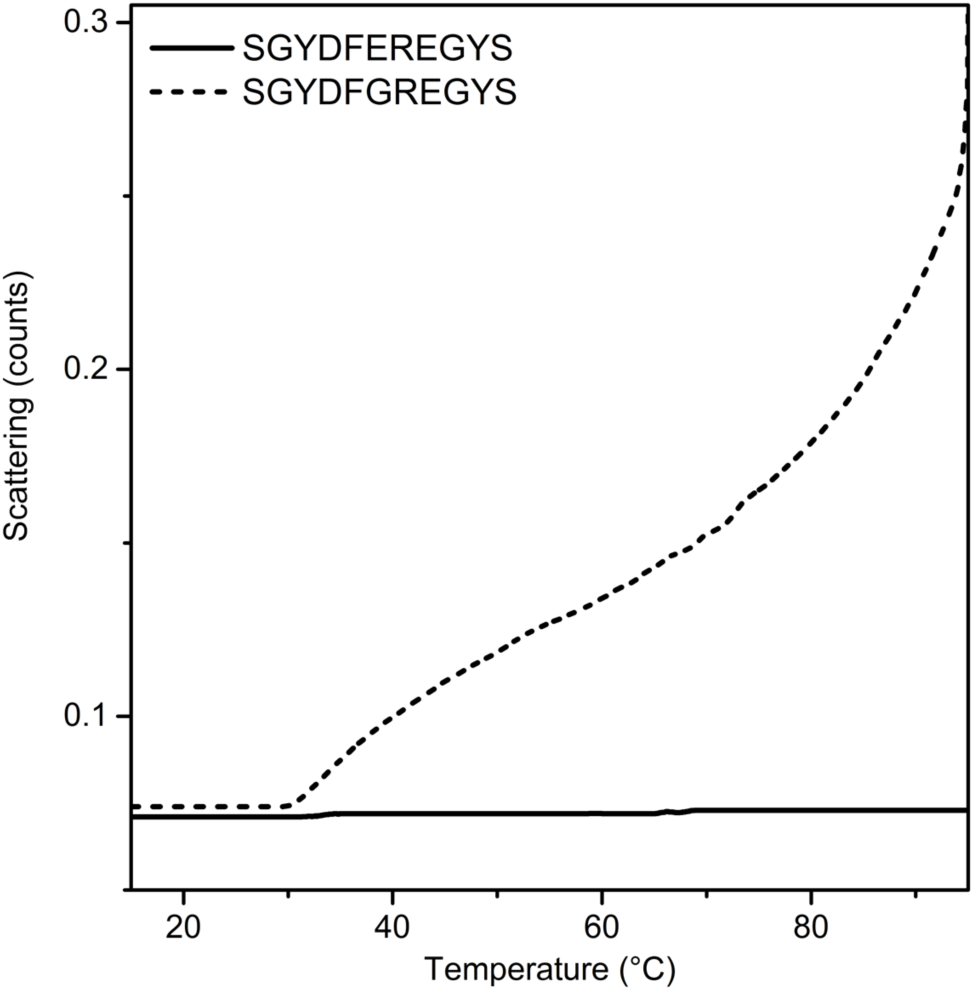
Temperature dependence of 540 nm absorption of peptide solutions at a concentration of 0.7 mg/ml.

In other words, the peptide corresponding to the wild-type sequence portion showed a tendency to self-association at temperatures above 30 degrees, while the one carrying the substitution did not aggregate. The presence of temperature-dependent oligomerization ability in the wt model peptide likely indicates a role of the glutamic acid residue at position 292 in self-association. In the peptide model, the presence of a substitution at the interface between the chains significantly affects: (i) the ability to interact with a protein containing an identical primary structure fragment; and (ii) the propensity for temperature-dependent self-association. These facts indirectly indicate a role of the glutamic acid residue at position 292 in the interface interaction during formation of the nucleoprotein complex.

To investigate the effect of E292G in temperature-dependent ribonucleoprotein complex structural changes, we performed small angle neutron scattering (SANS) measurements of the structures of RNP-containing wild-type and mutant protein (Figure 9).

**Figure 9.**
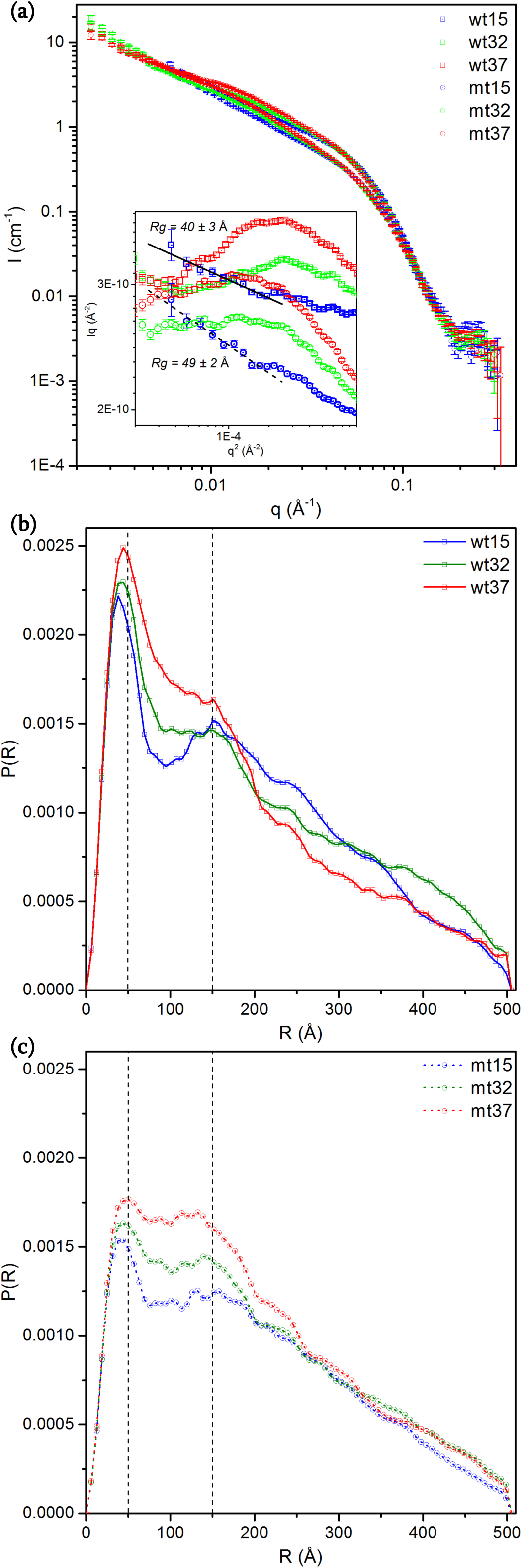
Small angle neutron scattering of wild-type (wt) and E292G mutant (mt) RNP solutions at 15°C, 32°C, and 37°C. (a) Representation in double logarithmic scale *I* vs *q*, where *I* – scattering intensity, *q* – magnitude of the momentum transfer; inset – Guinier coordinates (*Iq* vs *q*^*2*^). Distance distribution function *P(R)*, calculated for SANS spectra of wild-type RNP (b) and E292G mutant (c); dash lines mark *R* = 50 and 150 Å.

SANS shows that, at three temperatures (15°C, 32°C, 37°C), RNP structures form from both wt and mutant proteins leading to different spectra. At 15°C, the SANS data can be interpreted using a model featuring strongly elongated particles with a cross-sectional gyration radius of about 40 Å for the wt and of 50 Å for the RNP formed from substitution-carrying protein (Figure 9a, inset). These results match the published RNP double helix structure from the H1N1 influenza virus [5]. Temperature increases (from 15°C, to 32°C, and to 37°C) led to significant changes in SANS spectra, and while the behavior of the curves at low q was still close to the 1/q, noticeable deviations from the Guinier approximation for rod-like particles at small angles were observed, accompanied by an increase in the scattering of the samples in the q range between 0.01 and 0.03 Å^−1^. (Figure 9a, inset).

The distance distribution functions calculated from the experimental SANS spectra of RNP (Figure 9 b and c) show two distinct maxima at 4-5 nm and 15 nm. We have performed an analysis of the mean distance distribution between the centers of mass of all the monomer pairs in the model filament, averaged over the MD trajectory. Figure 10 shows the results of that calculation, with the pairs of monomers which would contribute to the experimentally observed peaks highlighted in the bottom-right half of each panel. Distances of approx. 5 nm were observed between the adjacent monomers along the same protein chain and also between the interacting monomers of the two filaments in the double helix. The distance of 15 nm corresponds to the pairs of monomers separated by one turn of the helix (Figure 11). It should be noted that, unlike the reference double-stranded helical nucleoprotein complex structure [5], in the MD simulated structure: both pair-distance measurement types (cis – on monomers of the same nucleoprotein complex strand; trans – on different strands, separated by 1 turn in the nucleoprotein complex helix) fall into the range 14-16 nm, thus giving rise to a peak in the distance distribution function [5].

**Figure 10.**
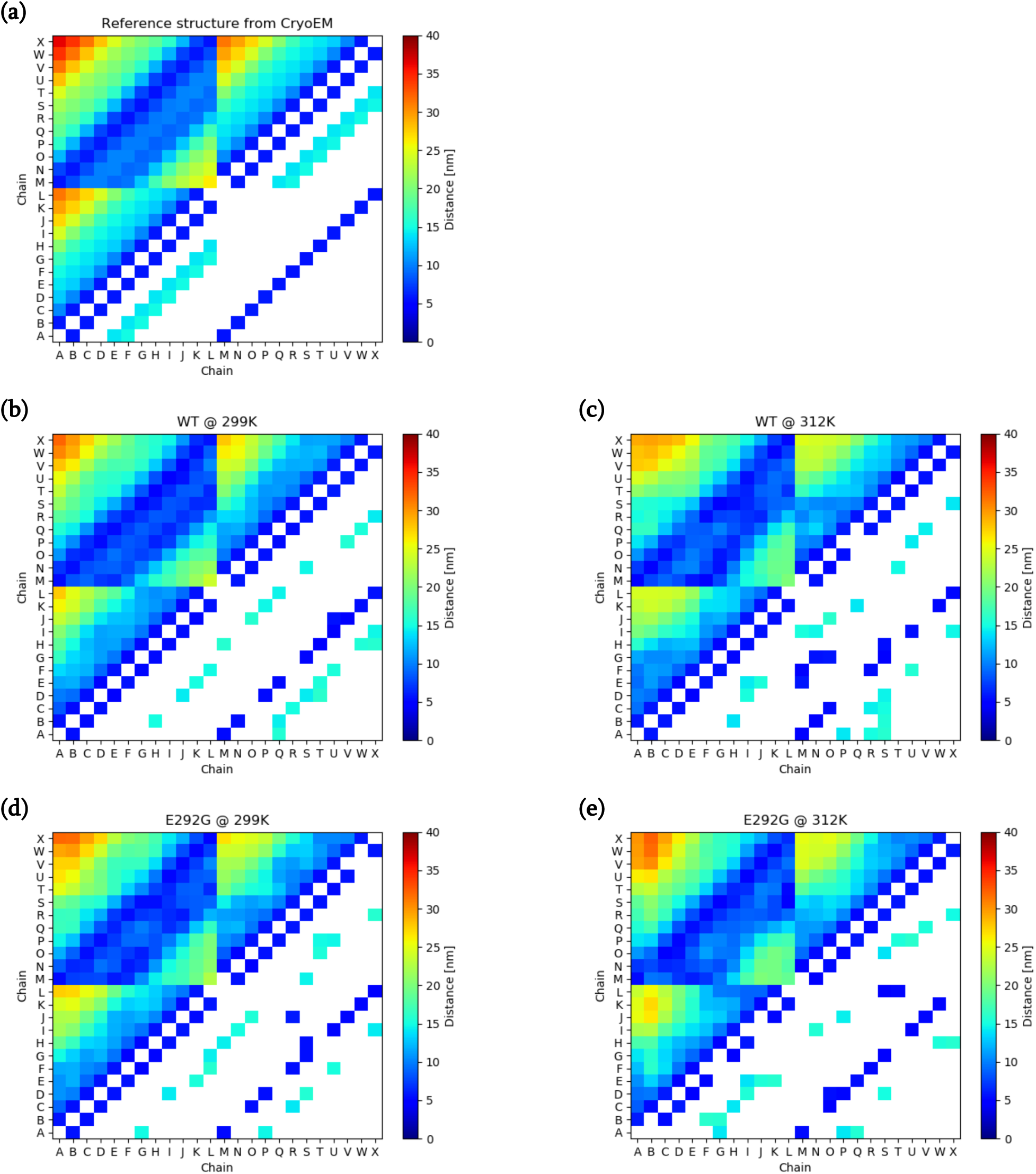
Mean distance distribution between the centers of mass of all monomer pairs, calculated for the reference structure [5] (a), for wild-type (wt), and for E292G mutant (mt) NP at 26°C (299K) and 39°C (312K) (b-e). “Chains” from A to L correspond to NP monomers in the first helix chain of RNP; those from M to X correspond to the second

**Figure 11.**
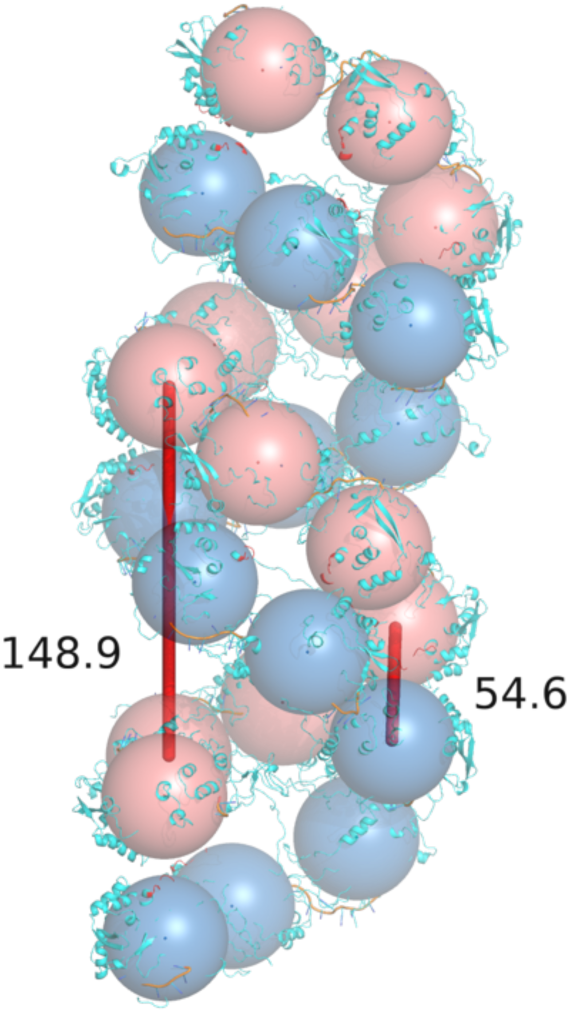
Model of RNP double-helix filament. Blue and red balls represent NP monomers. The characteristic distances between the monomers’ centers of mass (148.9 and 54.6 Å) are indicated by red bars.

The temperature-dependent differences in SANS of wild type and mutant RNP highlight structural differences between them. The mutant protein appears to form fewer contacts between potentially interacting regions of adjacent NP filaments. Loss of RNP’s rod-like shape at elevated temperatures, accompanied by a shift of the first P(R) peak toward larger distances (from 4 nm to approx. 5 nm), may be attributable to decreased interaction between regions in two adjacent NP filaments, leading to large distortions in RNP structure.

## Conclusions

Molecular dynamics simulation data, together with DSF and SANS results, suggest high structural lability of the NP filament; this likely permits partial unwinding of portions of the protein double helix near functional temperatures, a necessary step for RNA polymerase function. Using a peptide model, the region responsible for the strand interaction has been confirmed to contain E292 the residue. Apparently, the presence of amino acid substitution E292G, which is associated with cold adaptation, weakens interactions between NP strands in RNPs, thereby making possible their partial dissociation at lower temperature.

## Acknowledgments

This work was supported by Russian Science Foundation grant № 19-74-20146. The SANS experiments were performed on KWS-1 (proposal number 7879) and KWS-2 (proposal number 9634). The Prometheus PR.48 equipment used for nanoDSF experiments was provided by Nanotemper Technologies RUS LLC.

